# Assessing white matter pathway reproducibility from human whole-brain tractography clustering

**DOI:** 10.1101/833095

**Authors:** Jason Kai, Ali R. Khan

**Affiliations:** Department of Medical Biophysics, Schulich School of Medicine & Dentistry, The University of Western Ontario, London, Ontario, Canada; Centre for Functional and Metabolic Mapping, Robarts Research Institute, The University of Western Ontario, London, Ontario, Canada

## Abstract

Diffusion MRI, together with tractography techniques, is a non-invasive tool to investigate the brain’s structural pathways (tracts). These tracts join together different regions of the brain and tract identification often involves the use of manual ROIs or automated techniques such as clustering. By studying these connections, current understanding of the connectome can be improved and changes due to disease in patient populations may be identified. We developed a tool to automatically identify all pathways in the human brain, including the short-range, U-shaped tracts, and map quantitative scalar metrics along the pathway trajectory for subsequent analysis. Pathways are identified via a spectral clustering technique on two datasets: Human Connectome Project (intersubject) and MyConnectome Project (intrasubject) and the reliability of the extracted tract and scalar values are evaluated. Average Euclidean distances and volumetric overlap were computed and indicated good spatial reliability. Intraclass correlations of the fractional anisotropy value mapped along the tract was calculated and exhibited good reproducibility within each dataset. Additionally, these evaluation metrics, together with the coefficient of variation of the mean streamline count is used to determine reliably identified U-shaped tracts across the datasets. The developed tract identification tool is an additional resource to studying the human connectome with increased confidence in the results. The identified reliable U-shaped tracts contributes to the identification of common structural connections across individuals and aids in advancing our understanding of the brain’s short-range pathways.

## Introduction

The brain comprises of numerous regions connected by bundles of axons which form structural pathways [1, 2]. These pathways form networks to distribute information between different regions and enable performance of complex tasks [3]. Diffusion magnetic resonance imaging (dMRI) and tractography techniques allow for estimation of pathway trajectories. Due to the nature of axonal configuration, water molecules preferentially diffuses along the axis of an axon [4]. dMRI acquires information sensitive to the diffusion, enabling non-invasive investigation of these structural pathways [5]. Advancements in hardware and software have enabled the ability to study both long and short range (U-shaped) pathways, extending to distant regions of the brain and joining adjacent brain regions residing just below the cortical surface respectively [6, 7].

Tractography techniques are able to leverage the diffusion information to reconstruct pathway trajectories as streamlines [1]. A number of tractography algorithms have been developed and can be broadly categorized into one of two groups: deterministic or probabilistic. Regardless of the chosen algorithm, the tracking process is similar: (1) estimate fiber orientation within the imaging voxel and (2) follow along a fiber orientation to generate streamlines indicative of the estimated trajectory. The strengths and weaknesses of each type of algorithm have been previously reviewed by Sotiropoulos and Zalewsky [1] and Sarwar [2].

While the gold standard for investigating structural connectivity is the use of chemical tracers, tractography offer non-invasive methods to investigate and accurately reconstruct structural pathways [1, 3]. Past studies have implicated both the short-range [6,8,9] and long-range [10–12] pathways in patients with either neurological or psychiatric disorders. Improved understanding of how structural connections are affected may increase efficacy of treatment. Additionally, abnormalities affecting the axonal bundles could provide key insights to diagnosing neurological or psychiatric disorders. Previous studies have observed quantitative differences in white matter tracts (pathways joining together two brain regions) between healthy individuals and patients, and correlating observations with patients’ clinical symptoms [10].

To identify different tracts from tractography, manual or automated techniques can be employed. Manual techniques require users to place inclusion and exclusion regions of interest to extract tracts for further investigation. These techniques are often laborious, requiring knowledge of anatomy, and results can vary between different users or sessions [13]. In contrast, automated techniques employ unsupervised clustering algorithms to identify tracts [13] and have been previously shown to produce pathways with high confidence [13, 14]. These techniques often rely on the similarity of streamline trajectories. While automated techniques may include incomplete or false positive streamline generated from tractography [13], user biases introduced with manual intervention are avoided [14]. A number of automated techniques have been developed and applied to tractography (O’Donnell and Westin [15], Maddah *et al.* [16], Li *et al.* [17], Garyfallidis *et al.* [18] to name a few).

Reproducibility using any technique is crucial to increasing confidence of the results. With numerous algorithms available for performing tractography and its noted challenges [19, 20], it is important for clustering techniques to be able to replicate identified tracts with high reliability. Additionally, associated quantitative measurements along tract trajectories should be reproducible across different subjects or sessions [21]. In this work, we introduce our tractography clustering tool, which identifies both long and short range tracts, and evaluate the reproducibility of the outputs using this developed resource by applying it to two open source datasets: (1) Human Connectome Project, and (2) MyConnectome Project. Additionally, we aim to identify reliable U-shaped tracts using these datasets.

## Materials and methods

All processing is performed on a high performance compute cluster hosted by Compute Canada with Intel Xeon E5-2683 v4 processors (2.1 GHz) and 128GB of RAM using a containerized computing environment. Template creation and tractography generation pipelines were created via Nipype [22], a tool for creating reproducible pipelines written in Python. Code and container recipes for these pipelines can be found at the following Github repository: www.github.com/khanlab/mrtpipelines. The clustering tool used to identify structural pathways can be found at the following Github repository: www.github.com/khanlab/neurobeer.

### Data acquisition and pre-processing

#### Human Connectome Project

Pre-processed dMRI data of 100 unrelated subjects (46 male, 54 female) from the HCP1200 release of the Human Connectome Project (HCP) [23] was used to create a population template. An additional subset of 15 subjects (8 male, 7 female) was used to assess the intersubject reproducibility. dMRI data was acquired on a customized Siemens Skyra 3T scanner [24, 25] with the following scanning parameters: repetition time/echo time (TR/TE) = 5520 / 89.50 ms; resolution = 1.25 × 1.25 × 1.25 mm^3^; b-values = 1000, 2000, 3000 s/mm^2^ (90 directions each) with 18 b-value = 0 s/mm^2^ images. Full details regarding the acquisition of diffusion data can be found in the HCP 1200 subject reference manual (https://humanconnectome.org/study/hcp-young-adult/document/1200-subjects-data-release). Pre-processing of acquired data is performed as described in Glasser *et al.* [26].

#### MyConnectome Project

Multiple sessions of dMRI data of a single male subject acquired over a 3 year period as a part of the MyConnectome Project [27] was used to assess reproducibility of intrasubject data with our developed tool. Data acquisition was performed on a separate Siemens Skyra 3T scanner. 16 out of the 94 available production sessions from the project contained dMRI acquisitions. Scanning parameters are as follows: repetition time/echo time (TR/TE) = 5000 / 108 ms; resolution = 1.74 × 1.74 × 1.7 mm^3^; b-values = 1000, 2000 s/mm^2^ (30 directions each) with 4 b-value = 0 s/mm^2^ images. Detailed diffusion acquisition information can be found in the MyConnectome study protocol (http://myconnectome.org/wp/53-2).

Pre-processing of dMRI acquisitions was performed using an in-house pipeline. First, principal component analysis based denoising [28, 29] is performed followed by unringing of the dMRI data to correct for the effects of Gibbs ringing [30]. Afterwards, FSL’s topup [31, 32] and eddy [33] were applied to correct for distortions induced by susceptibility, eddy currents, and subject motion.

### Fiber orientation distribution

A fiber orientation distribution (FOD) template was created using the subset of 100 unrelated subjects of the HCP1200 release using the Mrtrix3 software suite [34]. First, a response function was estimated from each subject’s dMRI acquisition using the Dhollander algorithm [35] for the purposes of spherical deconvolution. This algorithm provides an estimate of the response for each tissue component: white matter (WM), gray matter (GM), and cerebrospinal fluid (CSF). An average response function for each tissue component was then computed from all the subjects’ response functions.

Using the average response function of each tissue component, FODs for each subject was individually computed via a multi-shell, multi-tissue spherical deconvolution algorithm [36]. Application of this algorithm computes a FOD estimate based on the unique b-value dependency of each component. Tissue component FOD estimates are normalised via multi-tissue informed log-domain intensity normalisation, correcting for both intensity differences and bias fields.

Following normalisation, an unbiased group-average FOD template was created. Creation of the population template was optimized with 3 registration steps (rigid, affine, and non-linear), with 6 iterations each in the rigid and affine registration and 15 iterations in the non-linear registration. Registration was performed following a multi-resolution pyramid and using Mrtrix3 default parameter settings. The template creation registers and reorients the FODs of each subject to the created template space. An average (population) template of fiber orientations was created from the registered and reoriented FODs [37]. The top row of Fig 1A exhibits the steps taken to produce the FOD population template.

**Fig 1.**
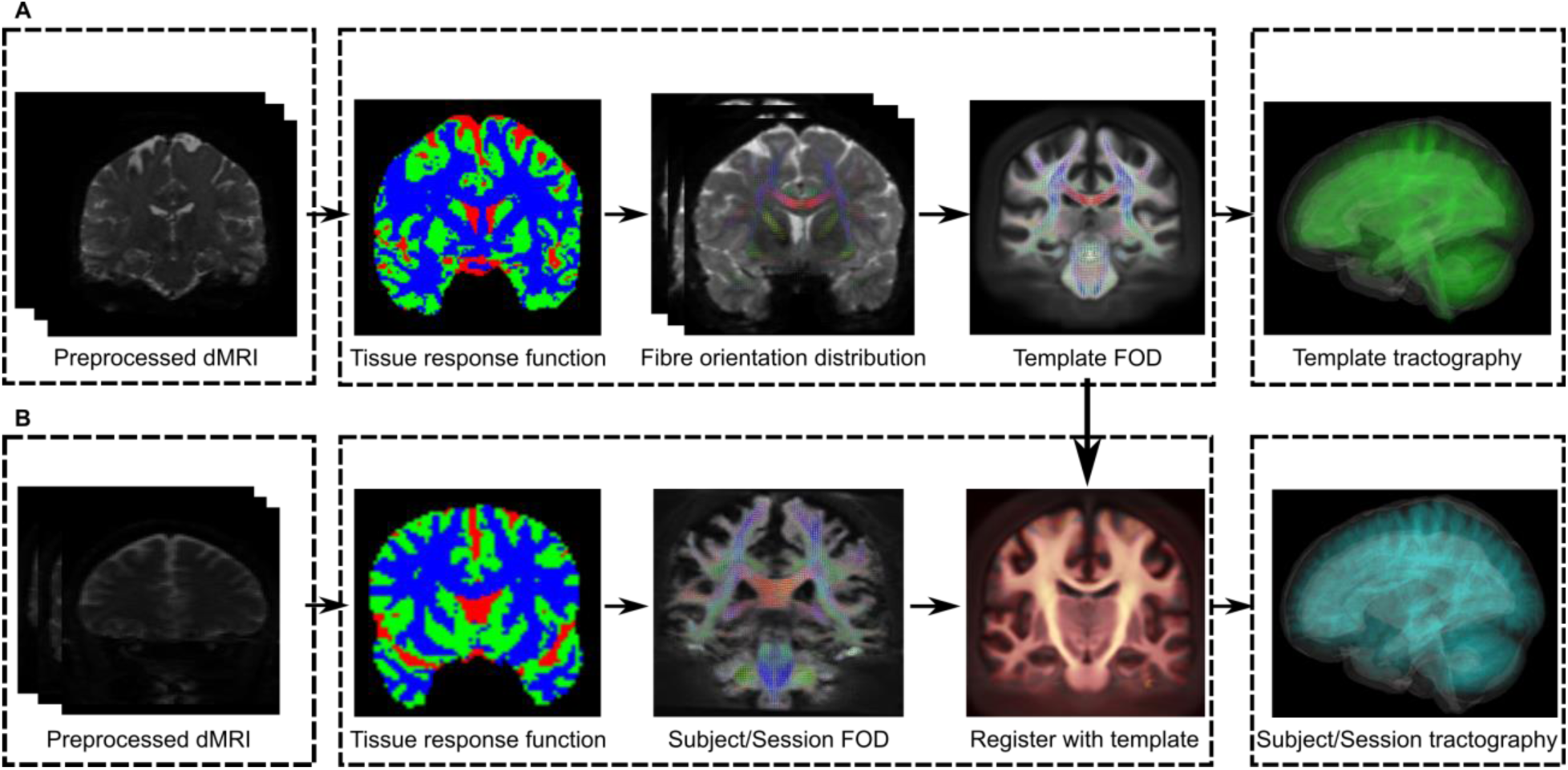
Processing pipeline for template creation and subject/session processing. (A) Minimal preprocessing of subject dMRI from HCP data to create population fibre orientation distribution (FOD) template and whole-brain probabilistic tractography. (B) Preprocessing steps taken for subject/session dMRI with in-house pipeline. Computed FODs were registered to template space followed by generation of whole-brain probabilistic tractography.

Similar steps are taken in computing FODs for session data from the MyConnectome Project. For each session, a response function for each tissue component was estimated using the Dhollander algorithm, however estimated response functions are not averaged. From the computed response function, FOD estimation was again performed with multi-shell, multi-tissue constrained spherical deconvolution, followed by FOD normalisation. Normalised FODs were then aligned to the created FOD template with FODs reoriented in the template space. Fig 1B displays the processing pipeline described in this section.

### Streamline tracking and quantification

Whole-brain probabilistic tractography was generated on the created template, MyConnectome sessions, and the 15 additional subjects from the HCP using Mrtrix3. Tracking of streamlines was performed using the iFOD2 algorithm [38], a second-order integration over FODs. This algorithm makes use of the previously computed FODs to identify streamlines such that the streamline has a higher probability of traversing a path where the amplitudes of FODs along the path are large or above the default FOD threshold of 0.05. Seeding of tractography was performed at random within the brain until a target of 100,000 streamlines have been selected.

Streamline densities from the generated whole-brain tractography were then filtered to match the FOD lobe amplitude until filtering has reached a user-defined streamline count. For the template, the user-defined streamline count was set to 50,000, while the streamline count for all other generated tractography was set to 100,000. Filtering was performed by making use of the implemented spherical-deconvolution informed filtering (SIFT) of tractograms in Mrtrix3 [39]. Lastly, the SIFTed tractography was converted to visualization toolkit (VTK) polydata format for additional quantification and clustering.

For reproducibility assessment, diffusion tensor measurements, such as fractional anisotropy (FA), were computed. To fit tensors, acquired diffusion weighted images (DWI) from each session were normalised and transformed into template space. Diffusion tensor images (DTI) were then estimated using an iteratively reweighted linear least squares estimation implemented in Mrtrix3 [40]. From the tensor images, quantitative measurements of FA, mean diffusivity (MD), radial diffusivity (RD), and axial diffusivity (AD) were computed [41, 42] and mapped to corresponding streamlines, enabling further quantitative analysis following clustering.

### Clustering

Bundles of streamlines from SIFTed whole-brain tractography were identified via spectral clustering techniques [43] implemented in our developed tool. Initial identification of tracts were performed on the created whole-brain tractography template followed by propagation of cluster labels to the data used for assessing reproducibility based on streamline similarity.

Similarity of individual streamlines from the template tractography were assessed by comparing with all other streamlines within the template. First, 20 equally-spaced samples along the length of streamline were taken. A minimum average, direct-flip distance, as described by Garyfallidis *et al.* [18], was computed between corresponding samples across streamlines and summed. An example of this is shown in Fig 2A. Individual streamlines, where the distance was more than two standard deviations from the average whole-brain streamline distance, were discarded. A Gaussian kernel, with a width (*<u>σ</u>*) of 8mm, was applied to the computed distances to create an affinity matrix characterizing the similarity between streamlines for spectral clustering.

**Fig 2.**
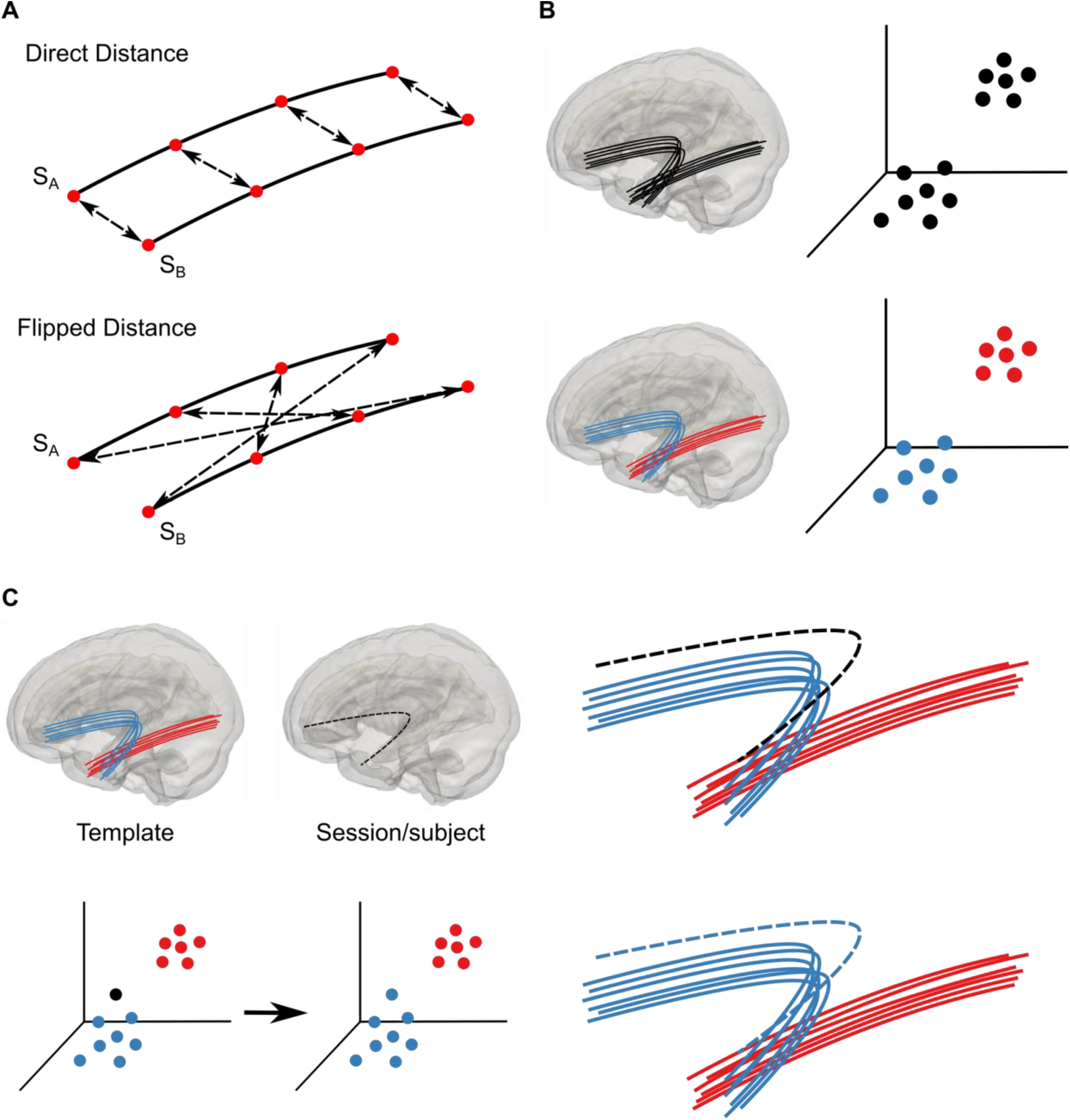
Spectral clustering steps to compare and group streamlines. (A) Equally-spaced samples are taken to assess trajectory similarity of streamlines with a minimum, average direct-flipped distance metric. Streamlines that are more similar exhibit shorter average Euclidean distances. (B) A simplified example of label assignment of two bundles of streamlines. Streamlines are represented as single, unlabeled points in higher dimensional spectral space (shown here in 3D). Labels are assigned after applying the k-means clustering algorithm in spectral space. (C) Propagation of cluster label to a new session/subject streamline (dashed line) by comparing with labelled template streamlines (solid line). Label of most similar (nearest cluster) is assigned to the new streamline.

As described by Von Luxburg [43], Laplacian matrices are one of the primary tools used in spectral clustering. The spectral clustering implemented with our tool closely follows that as described by Ng *et al.* [44] and makes use of normalized Laplacian matrices. Generalized eigenvectors and eigenvalues were computed and arranged in ascending eigenvalue order from the normalized Laplacian matrices. The first k eigenvectors were selected as the embedding vectors to be used in the spectral space and is determined by identifying the two consecutive eigenvalues with the greatest difference (gap statistic), resulting in selection of stable embedding vectors. Next, the number of clusters to be identified was specified and a total of 800 clusters was chosen for clustering of the template whole-brain tractography. The individual streamlines were embedded into spectral space as single points using the spectral clustering technique and the k-means algorithm was applied (where k = 800) to label individual streamlines and assign each to an appropriate cluster. Established clusters were coloured according to the coordinates of the cluster centroids, as described in Brun *et al.* [45]. An example of clustering similar streamlines is seen in Fig 2B.

Labels were propagated to the whole-brain tractography generated for individual subjects/sessions assessed for reproducibility. A sub-sample of the labelled template tractography was extracted to improve memory consumption and processing time. Similarity of streamlines between the sub-sampled template and individual subject/session tractography was computed with the previously described method. Again, a Gaussian kernel of width (8mm) was applied following outlier removal. Labels from template streamlines were then propagated to subject/session tractography based on greatest similarity. This establishes label correspondence between the most similar streamlines across generated tractography. Shown in Fig 2C, presents the workflow required to propagate labels from template to new subject/sessions.

### Short range, U-shaped streamlines

Streamlines comprising short-range, U-shaped tracts were identified using adapted parameters [46, 47] and implemented as a filter to extract from whole-brain tractography. The identified U-shaped streamlines can be clustered using the same method as explained in the previous section, replacing whole-brain tractography with the filtered, U-shaped tractography. Extraction of U-shaped streamlines take advantage of the shape and curvature by leveraging the distance between the endpoints of a streamline and the streamline length. Streamline lengths were computed as the arc length (L) of the sample points (s_i_). Endpoint distances (D) were computed as the Euclidean distance between each terminal end of the streamline.

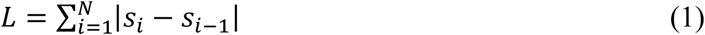

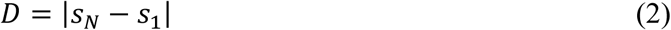

To extract only streamlines with an U-shaped curvature, the endpoint distance was constrained to approximately one-third of the streamline length. Additional length constraints of 20mm (minimum) and 80mm (maximum) were imposed. Streamlines crossing the mid-sagittal plane of the brain (eg. corpus callosum) were removed. Fig 3 demonstrates the features examined to identify U-shaped streamlines from whole brain tractography.

**Fig 3.**
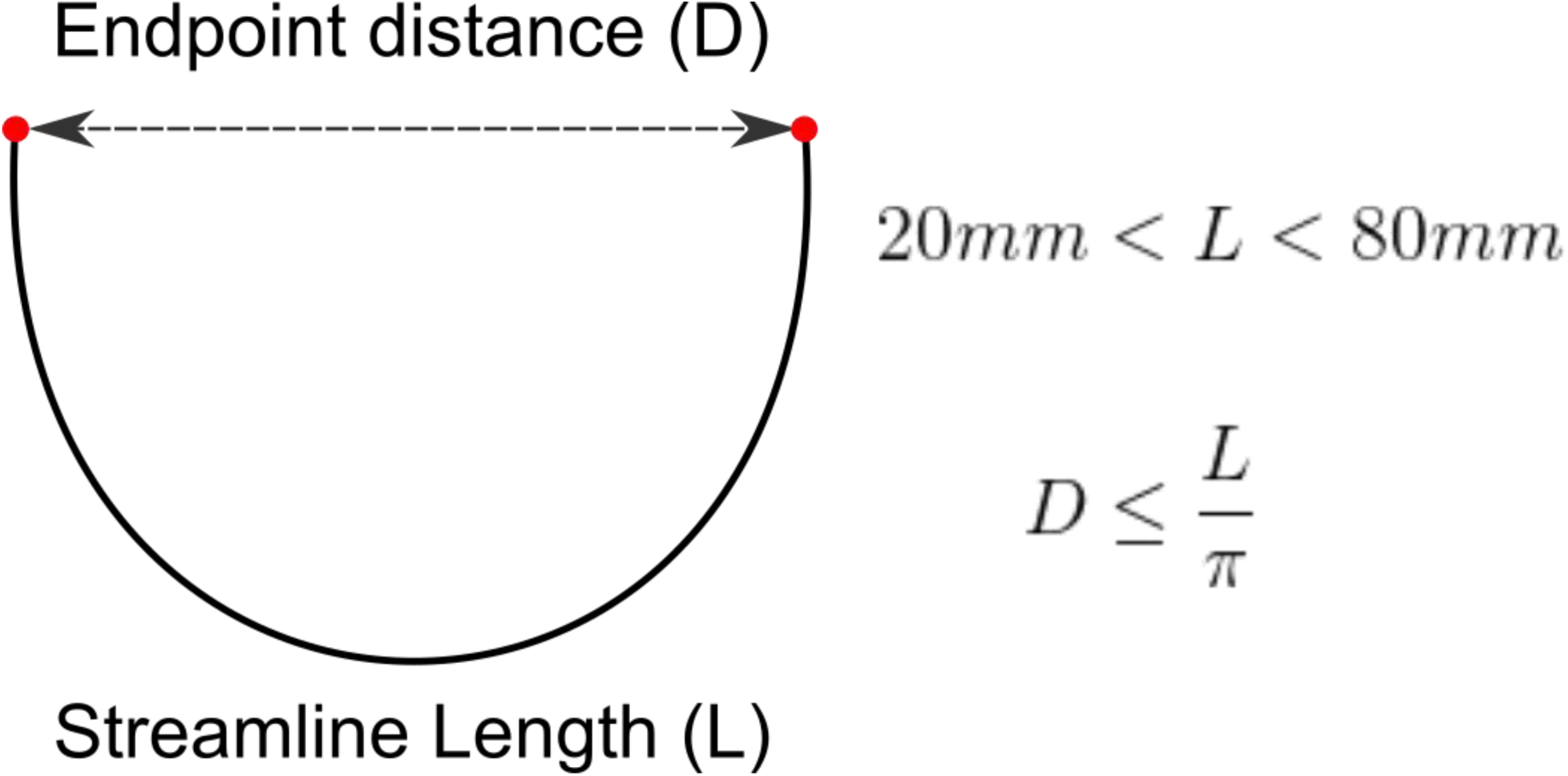
Identification of short-ranged, U-shaped streamline. Comparison of streamline line length (L) and endpoint distance (D) is performed to determine if streamline trajectory is U-shaped. Endpoint distance is constrained to approximately one-third of streamline length. Additional minimum and maximum length constraints are imposed (20mm and 80mm respectively).

## Results

A total of 15 tracts (5 in each brain hemispheres and along the corpus callosum) were identified for evaluation of reproducibility using the template. For hemispheric tracts, the corresponding tract in the opposite hemisphere was selected for evaluation.

### Average Euclidean distance

For each identified tract, a centroid streamline was computed (Fig 4A) and the average Euclidean distance was computed from corresponding sample points between the centroid streamlines of a tract for a single subject/session and of the template tract (Fig 4B). The majority of identified tracts fell within 4mm (or 4 voxels) of the template centroid streamline. The average Euclidean distance observed across investigated tracts was 3.15 +/− 0.394 mm and 3.28 +/− 0.652 mm across sessions (intrasubject) and subjects (intersubject) data respectively (Fig 4C). A notable exception was observed in the right optic radiation (R OR) where the average Euclidean distance was 8.07 +/− 0.271 mm and 7.93 +/− 0.845 mm for the intrasubject and intersubject data respectively.

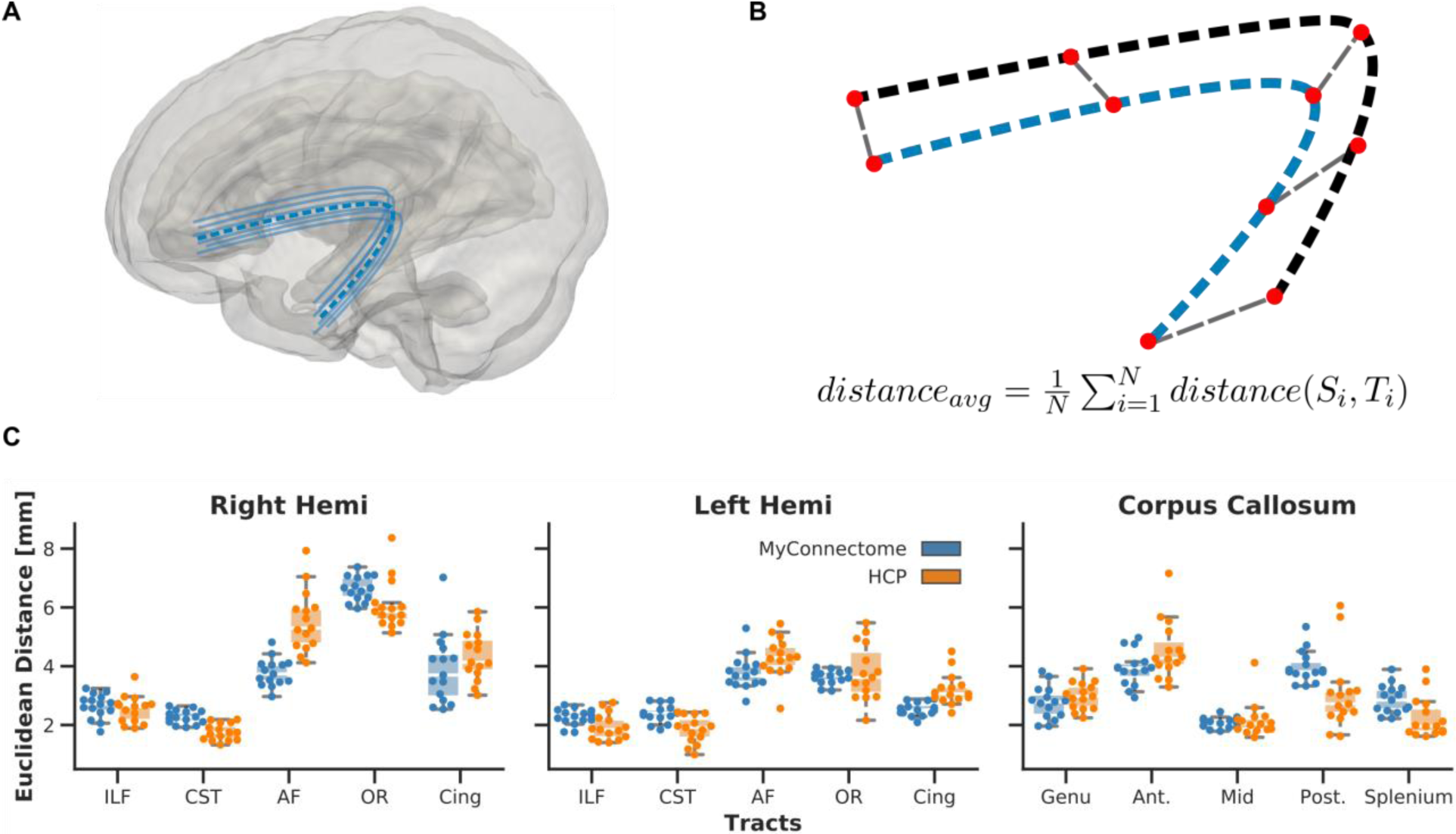
(A) Centroid streamline (dashed line) within a bundle is identified (B) Average distances between corresponding sample points are computed between subject/session streamline (blue dashed line), S_i_, against template (black dashed line) centroid, T_i_. (C) Computed average Euclidean distance for each session (MyConnectome) or subject (HCP) is shown as a single point for each identified tract.

### Intersection over union

Voxel-based analysis was also performed to analyze the spatial similarity of the tracts. Voxels where streamlines passed through were identified, creating a binary map, and was spatially smoothed with a Gaussian kernel of 0.5mm width. From the smoothed map, the average intersection over union (IOU) was across computed, again comparing tracts identified within each group (intrasubject vs intersubject) to the template tract, and within the group. An example of the comparison performed can be seen in Fig 5.

**Fig 5.**
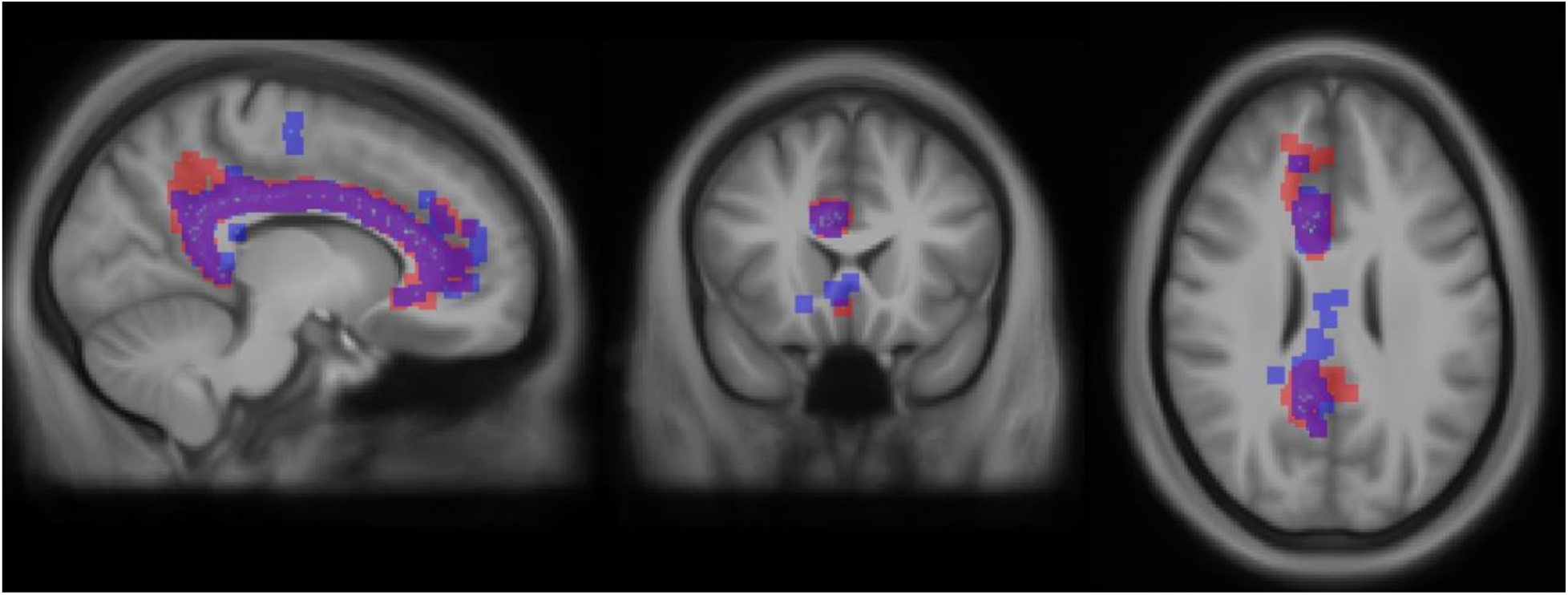
Example of intersection over union of right cingulum (R Cing) Volume of tract identified in a MyConnectome session (red) and the created template (blue) is overlaid atop each other to visualize the intersection of the two tracts. Similar comparisons were performed for all identified tracts and averaged.

A summary statistic for each tract within a group can be found in Table 1.

**Table 1.** Intersection over union summary.

| Cluster | Template |  | Within group |  |
| --- | --- | --- | --- | --- |
|  | MyConnectome | HCP | MyConnectome | HCP |
| R ILF | $0.471 \pm 0.0137$ | $0.451 \pm 0.0157$ | $0.639 \pm 0.137$ | $0.626 \pm 0.143$ |
| R CST | $0.369 \pm 0.0119$ | $0.436 \pm 0.0120$ | $0.681 \pm 0.121$ | $0.648 \pm 0.135$ |
| R AF | $0.384 \pm 0.0134$ | $0.353 \pm 0.0305$ | $0.598 \pm 0.153$ | $0.484 \pm 0.204$ |
| R OR | $0.384 \pm 0.0144$ | $0.395 \pm 0.0253$ | $0.641 \pm 0.136$ | $0.590 \pm 0.156$ |
| R Cing | $0.430 \pm 0.0144$ | $0.449 \pm 0.0168$ | $0.629 \pm 0.142$ | $0.628 \pm 0.144$ |
| L ILF | $0.467 \pm 0.00863$ | $0.448 \pm 0.0122$ | $0.652 \pm 0.133$ | $0.638 \pm 0.138$ |
| L CST | $0.446 \pm 0.0127$ | $0.478 \pm 0.0122$ | $0.654 \pm 0.132$ | $0.625 \pm 0.143$ |
| L AF | $0.334 \pm 0.00977$ | $0.344 \pm 0.0136$ | $0.595 \pm 0.155$ | $0.546 \pm 0.176$ |
| L OR | $0.443 \pm 0.00637$ | $0.446 \pm 0.00980$ | $0.641 \pm 0.136$ | $0.613 \pm 0.147$ |
| L Cing | $0.377 \pm 0.00932$ | $0.381 \pm 0.0110$ | $0.626 \pm 0.143$ | $0.617 \pm 0.148$ |
| CC - Genu | $0.400 \pm 0.0155$ | $0.394 \pm 0.0125$ | $0.539 \pm 0.175$ | $0.509 \pm 0.186$ |
| CC - Body (Ant) | $0.391 \pm 0.0162$ | $0.378 \pm 0.0343$ | $0.524 \pm 0.181$ | $0.480 \pm 0.199$ |
| CC - Body (Mid) | $0.429 \pm 0.0200$ | $0.427 \pm 0.0236$ | $0.657 \pm 0.131$ | $0.605 \pm 0.153$ |
| CC - Body (Post) | $0.379 \pm 0.0151$ | $0.379 \pm 0.0271$ | $0.568 \pm 0.164$ | $0.514 \pm 0.187$ |
| CC - Splenium | $0.435 \pm 0.0132$ | $0.448 \pm 0.0211$ | $0.620 \pm 0.144$ | $0.590 \pm 0.156$ |

Intersection over unions (mean ± standard deviation) is computed for tracts identified in MyConnectome and HCP datasets. Comparisons are performed both against tracts identified from the template and within each dataset. Yellow and blue shaded cells indicate tracts identified in the right and left hemispheres respectively.

### Intraclass correlation

Additionally, along tract reproducibility of a quantitative scalar metric was evaluated. Here, fractional anisotropy (FA) is chosen as the metric subject to evaluation due to its widespread use and interpretation. To evaluate its reproducibility, the scalar value at each sample location was taken and grouped prior to computing the intraclass correlation, a measure of absolute agreement for the values within a group. In Fig 6, each data point represents the average FA value at a particular sample position for a particular tract. High intraclass correlation (ICC > 0.8) was observed in both intrasubject and intersubject groups indicating good agreement of FA values at each sample location, for all identified tracts. Intrasubject data generally exhibited greater agreement than intersubject data, with the left cingulum (L Cing) as the only exception.

**Fig 6.**
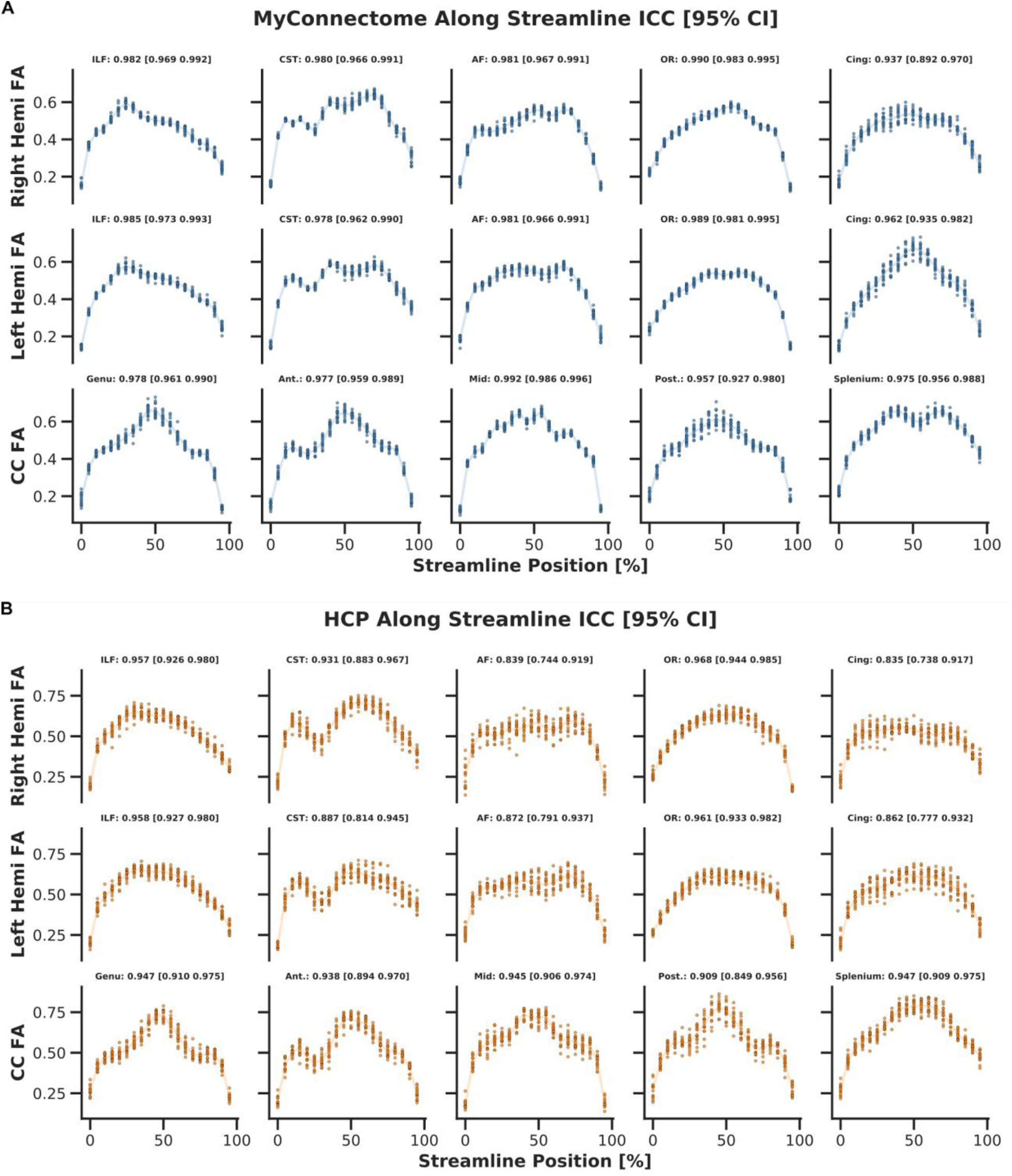
Along tract average fractional anisotropy. Average fractional anisotropy (FA) is computed for each tract in subject/session and grouped by sample location. Grouped data points are used to compute intraclass correlation ICC,(summarized in Table 2. (A) MyConnectome (intrasubject) ICC (B) HCP (intersubject) ICC

Table 2 summarizes the computed ICCs for a tract within each group.

**Table 2.** Along tract intraclass correlation of fractional anisotropy.

| Cluster | MyConnectome | HCP |
| --- | --- | --- |
| R ILF | 0.982 [0.969, 0.992] | 0.957 [0.926, 0.980] |
| R CST | 0.980 [0.966, 0.991] | 0.931 [0.883, 0.967] |
| R AF | 0.981 [0.967, 0.991] | 0.839 [0.744, 0.919] |
| R OR | 0.990 [0.983, 0.995] | 0.968 [0.944, 0.985] |
| R Cing | 0.937 [0.892, 0.970] | 0.835 [0.738, 0.917] |
| L ILF | 0.985 [0.973, 0.993] | 0.958 [0.927, 0.980] |
| L CST | 0.978 [0.962, 0.990] | 0.887 [0.814, 0.945] |
| L AF | 0.981 [0.966, 0.991] | 0.872 [0.791, 0.937] |
| L OR | 0.989 [0.981, 0.995] | 0.961 [0.933, 0.982] |
| L Cing | 0.962 [0.935, 0.982] | 0.862 [0.777, 0.932] |
| CC - Genu | 0.978 [0.961, 0.990] | 0.947 [0.910, 0.975] |
| CC - Body (Ant) | 0.977 [0.959, 0.989] | 0.938 [0.894, 0.970] |
| CC - Body (Mid) | 0.992 [0.986, 0.996] | 0.945 [0.906, 0.974] |
| CC - Body (Post) | 0.957 [0.927, 0.980] | 0.909 [0.849, 0.956] |
| CC - Splenium | 0.975 [0.956, 0.988] | 0.947 [0.909, 0.975] |

Computed intraclass correlation (ICC) [95% confidence interval] of fractional anisotropy for each identified tract. Yellow and blue shaded cells indicate tracts identified in right and left hemispheres of the brain respectively.

### Streamline count

Streamline counts for each identified tract within a subject / session was also explored. Fig 7 visualizes these streamline counts as a matrix. The left corticospinal tract (L CST) was consistently identified as one of the investigated tracts comprising more streamlines in both intrasubject and intersubject data. Additionally, it was noted the right arcuate fasciculus (R AF) and cingulum bundles of both hemispheres (L and R Cing) had the fewest number of streamlines amongst the investigated tracts. Findings regarding streamline counts were consistent across different scan sessions in the intrasubject data and the different subjects in the intersubject data. Differences in streamline counts were more pronounced in the intersubject data.

**Fig 7.**
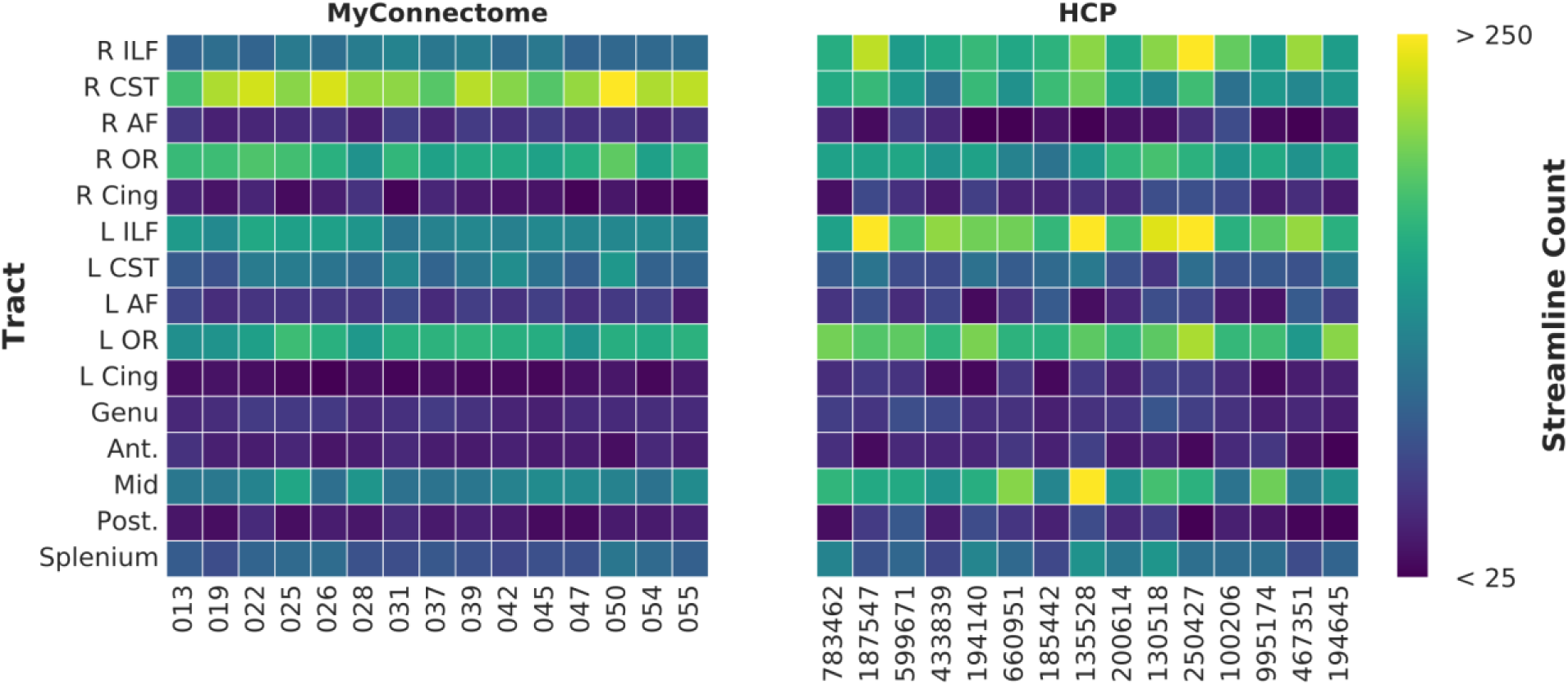
Streamline count matrix for each dataset. Matrix containing streamline counts for each tract (row) and subject/session (column) for MyConnectome (left) and HCP (right) tractography data.

### U-shaped streamlines and reliability

In assessing short-association tracts, it was observed that not all subjects/sessions contained streamlines in all tracts. Previously considered measures of average Euclidean distance between subject/session tract to template, intraclass correlation, and tract streamline counts can aid in the assessment of reliable short-association tracts across individuals. Fig 8 displays the average Euclidean distance, streamline counts and intraclass correlations in separate matrices for each dataset. Tracts identified in the sessions of the MyConnectome dataset where streamlines were available for all sessions displayed greater consistency in Euclidean distance and streamline count. Streamline counts showed less consistency between the different subjects of the HCP dataset. Importantly, the streamline count matrices enabled greater visualization of tracts lacking streamlines.

**Fig 8.**
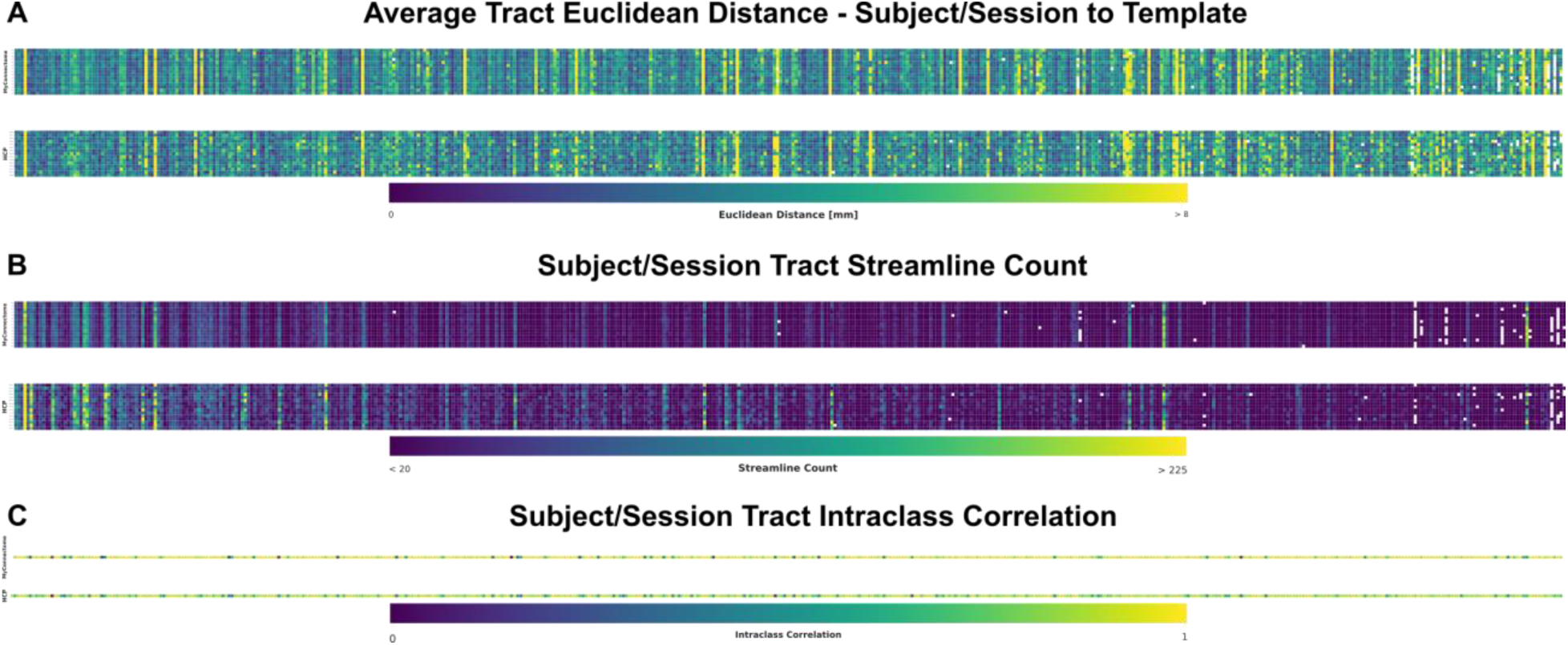
Matrices displaying metrics used for evaluating reliability of short-association U-shaped tracts. MyConnectome and HCP metrics are displayed in the top and bottom matrices respectively. Columns correspond to identified tracts. Tracts without streamlines for at least a single subject/session is displayed as a white box. (A) Average Euclidean distance between subject/session tract centroid to template centroid. (B) Streamline count for each subject/session tract. (C) Intraclass correlation for for each tract computed across corresponding sample points.

All U-shaped tracts with streamlines in all sessions/subjects in each dataset were sorted in descending coefficient of variation (CoV) (Fig 9A). CoV was not calculated for tracts missing streamlines in at least one session or subject. A combination of CoV and streamline counts were used as thresholds to identify reliable U-shaped tracts. Selected CoV thresholds of <20% and <40% were selected for MyConnectome and HCP datasets respectively. Additionally, a minimum and maximum mean streamline threshold of 10 and 150 respectively was applied. Fig 9B display the remaining U-shaped tracts in each anatomical view following thresholding.

**Fig 9.**
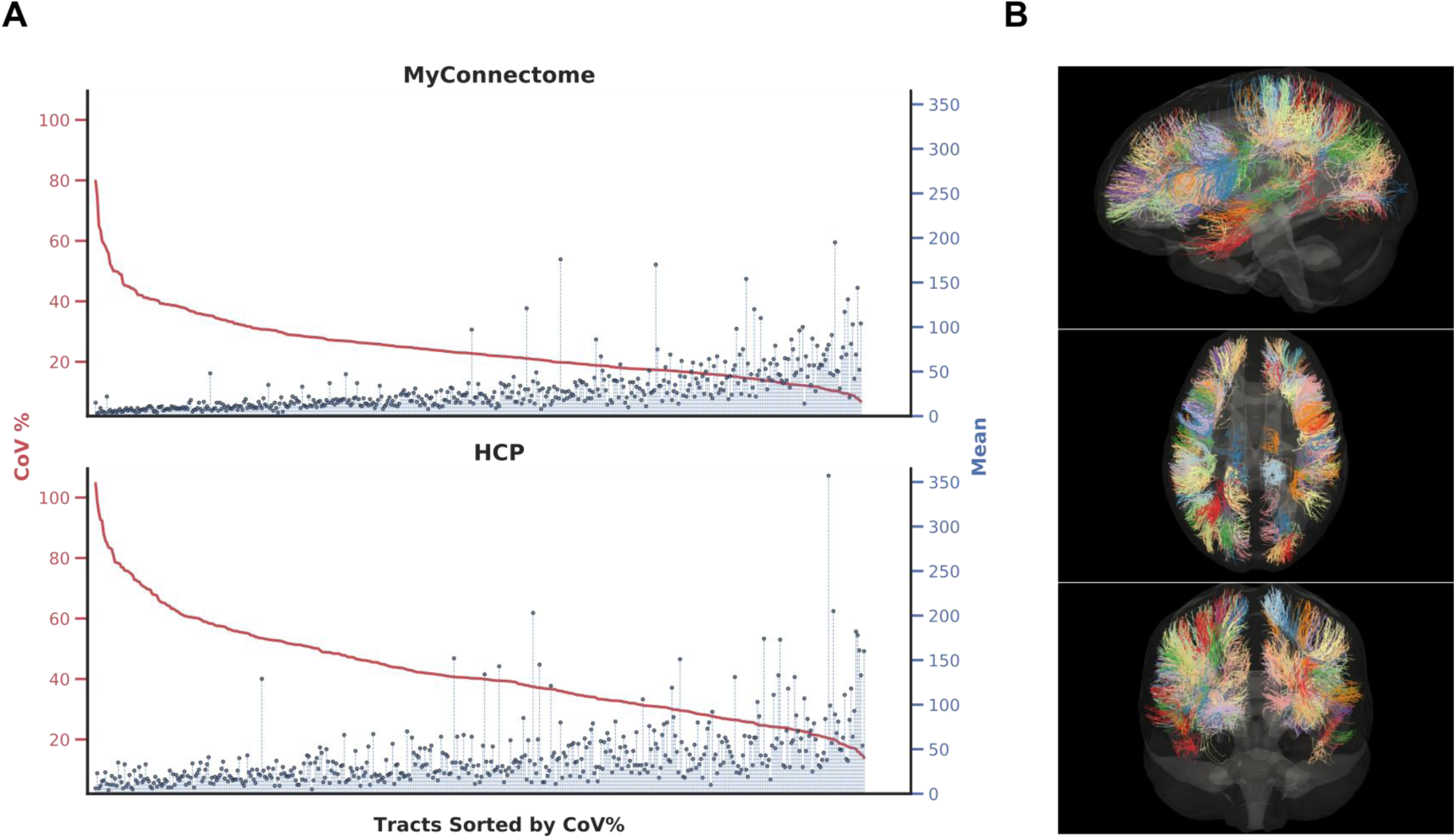
Identification of reliable short-range, U-shaped tracts. (A) Sorting of identified tracts for each dataset by coefficient of variation percentage (CoV %) in descending order. Tracts where streamlines were missing in at least a single subject/session are not shown. (B) Tracts determined to be reliably labelled in both datasets are identified through a combination of thresholding mean streamline counts and CoV% are displayed in axial (top), sagittal (middle), and coronal (bottom) views.

## Discussion

### Tractography assumptions

In order to perform clustering of streamlines and to compare scalar metrics associated with tracts across subjects and sessions, a few assumptions are made. As equally-spaced sample points were taken along streamlines, including the terminal ends, spatial correspondence between the sample points across the different streamlines was assumed. As a result, all streamlines are required to be sampled with the same number of points. However, streamlines that comprise a tract could be of different lengths, resulting in differences in spatial location of sample points. For example, an individual streamline may have already terminated prior to the terminus of longer streamline within the tract. As a result, the sample point located at the terminus end of the streamline may have better correspondence with a non-terminal sample point of the longer streamline. These differences in streamline lengths may affect the clustering of individual streamlines to tracts due to spatial differences of sample points. An additional consequence is the effect on evaluating of scalar values mapped along the streamlines of a tract. It would be expected that corresponding sample points within a tract should have similar quantitative measurements. However, if the sample points do not spatially correspond (as in the given example), quantitative measurements may vary due to extraction of values at different spatial locations.

Some studies have explored cutting off tractography streamlines prior to terminal ends [48, 49] such that streamlines within a bundle are of the same length. These studies typically retain the body of a tract, while eliminating the head and tail, where trajectories may be more varied. While these methods are able to better guarantee spatial correspondence of the samples taken along the tract, the extraction and subsequent evaluation of these structural pathways are not performed on the entirety of the tract.

### Evaluation metrics and uses

A number of metrics were used to evaluate reproducibility of identified tracts. Spatial reliability of identified tracts was determined by computing the mean Euclidean distance between the subject/session tract centroid and corresponding template tract centroid. The spatial reliability compared against the template is important to consider due to registration step in the generation of tractography and establishing corresponding clusters in each subject/session data. Reliable tracts should exhibit minimal distances when compared against the template tract. Exceptions were observed in the arcuate fasciculus and optic radiation residing in the right hemisphere (R AF and R OR respectively from Fig 4). In particular, the R AF in the intersubject dataset exhibited greater distance from the template tract. One possible explanation resides in the difficulty in tracking of the R AF. A previous study noted differences in the principal diffusion direction between subjects where the R AF could and could not be identified using deterministic tractography algorithms [50]. These differences may also affect probabilistic tracking algorithms, though not to the point where the R AF is lost completely. Additionally, neighbouring streamlines of other tracts with similar trajectories could be mislabelled and clustered with those of the R AF, such as the superior longitudinal fasciculus. The R OR in both datasets also demonstrated a large distance from the template tract. Upon visually inspecting this tract in the template, outlier streamlines that exhibited similar trajectories were seen in the identified template. These streamlines had terminal ends in similar spatial locations as the R OR, but with a more linear trajectory that was not removed during the outlier detection step. The manual removal of these streamlines resulted in a noticeable decrease in Euclidean distance.

Additionally, the intersection over union (IOU) was computed (results shown in Table 1), demonstrating the overlap of identified tracts with the template and also within each group of datasets. Within group IOU was greater than the IOU when compared against the template, with the intrasubject data exhibiting the greatest IOU in the majority of the tracts. Using the IOU, it was determined the volumetric agreement both against the template and within each group was moderate. It was noted in the binary maps that spurious end points resulted in poor overlap at these terminal locations. While there appeared to be good agreement within the body of the tracts, the poor overlap at the terminal ends may be a factor in the moderate agreement observed.

In order to assess the reliability of scalar metrics, intraclass correlation (ICC) of fractional anisotropy (FA) was computed and mapped along the tract. Comparisons were made within each group, with both groups exhibiting ICC values greater than 0.80, indicative of excellent reliability along the tract. The intrasubject dataset consistently demonstrated better reliability relative to the intersubject dataset in corresponding tracts as would be expected in single-versus multi-subject data. It is worth noting that tracts which were difficult to track (e.g. R AF) exhibited lesser reliability relative to the other investigated tracts amongst the intersubject dataset where anatomy may differ. Nonetheless, these results indicate excellent reproducibility of the scalar metrics in addition to the spatial reliability.

Lastly, for the evaluation and identification of reliable U-shaped tracts, the previously mentioned Euclidean distance and ICC metrics were evaluated in addition to streamline counts of each tract. Importantly, the coefficient of variation (CoV) percentage of the streamline count for each tract was computed and used (1) to assess the extent of the variability for each tract within a dataset, and (2) to determine an appropriate threshold for eliminating unreliable U-shape tracts. While registration can align different subject / session data, individual differences in local folding patterns of the brain could cause some streamlines to be considered outliers when using a template. As a result, some of the short-range tracts may not be identified. In spite of this, there are still common gyri and sulci amongst individuals. One of our aims was to identify reliable and discard unreliable U-shape tracts using our developed tool. Unreliable U-shape tracts was defined as those missing streamlines in a single subject/session within the dataset, average streamlines fewer than 10 (where certain session/subjects may only comprise of a few streamlines) or greater than 150 (due to inclusion of outlier streamlines or streamlines from other similar tracts). In total, 125 reliable U-shaped tracts were identified and over 350 tracts were discarded following the application of thresholds.

These results demonstrate the reliability of clustering methods both within subject (across multiple sessions) and across different subjects in identifying similar tracts and evaluating the scalar metrics along these tracts. Additionally, these results also present the challenge of evaluating the reliability of U-shaped tracts. Ultimately, these metrics can be used to identify tract reliability and improve or create new connectome atlases that include previously excluded tracts such as the U-shaped tracts joining local brain regions. Some progress has already been made on this front using atlas-based segmentations [47].

### Clustering vs regions of interest

Identification and extraction of tracts from whole-brain tractography can be performed a number of ways. In this work, tract identification was performed via clustering streamlines based on trajectory similarity. Other methods of tract identification make use of regions of interest (ROI) drawn manually or from an atlas, as inclusion or exclusion masks. ROI-based methods are able to identify streamlines joining together regions defined by inclusion masks. As noted previously, these methods are often time-consuming and rely on prior anatomical knowledge. Additionally, if an atlas is used to define ROIs, accurate registration with subject data is required.

Conversely, clustering methods do not require any prior knowledge. Instead a similarity metric is computed to characterize the affinity between streamlines. Clustering algorithms can then be applied to group to streamlines with high affinity. Registration between datasets is not explicitly required, though it is helpful in computing streamline similarity across datasets and to establish correspondence between different subjects. Automated techniques, such as clustering, ultimately reduce the amount of time required to identify the different tracts by eliminating the need to place inclusion / exclusion ROIs.

Clustering of whole-brain tractography is not without its own drawbacks. To compute similarity between streamlines, there must be the same number of sample points along the streamline. As mentioned previously, this could result in inaccurate comparison of streamlines due to differing lengths. Additionally, the number of clusters needs to be predefined by the user. There are ways to estimate the number of clusters (e.g. gap statistic, elbow method, etc.) that provide a starting value, though determining the optimal number is not straightforward. Initialization of clustering algorithms (e.g. k-means) are often random with final clusters dependent on the initialization of cluster centroids and the number of iterations the algorithm performs.

### Applications

Tract identification has a number of applications and implications both clinically and in research. As previously mentioned, one such application is the improvement or creation of atlases to include previously unknown or uncommon tracts are included (such as the U-shaped tracts). Identification of tracts can also aid in answering research questions pertaining to structural connectivity of the brain and for studying changes to such connections in patient populations or due to disease progression. Tractography can also play a role in surgical planning to either identify tracts to avoid or target tracts.

### Limitations

Whole-brain tractography generation and clustering is a computationally intensive task. Often, hundreds of thousands to millions of streamlines comprise a whole-brain tractogram resulting in large files. The amount of memory required to perform tasks, such as comparing and storing similarity of streamlines, is often large and increases with the number of streamlines that comprise the tractogram. Additionally, a large amount of processing time (at a minimum, a few hours for whole-brain tractography) is needed, again increasing with the number of streamlines.

Challenges associated with tractography have been well documented [51] and the issue of false positive streamlines was noted [20]. An attempt at addressing this issue was performed through the elimination of streamlines based on outlier detection. Further constraints could be employed to help clean up streamlines, such as the use of anatomically constrained tractography or quantitative information. While this may aid elimination of some false positive streamlines, it is acknowledged that not all false streamlines are removed nor are all identified clusters of streamlines valid (e.g. joining 2 brain regions that is not actually connected). Additionally, regions of streamline kissing or crossing can create problems tracking streamlines. To tackle this problem, constrained spherical deconvolution (CSD), probabilistic tracking and spherical-deconvolution informed filtering techniques are employed. Together, these methods have shown an improvement in tracking streamlines in the problematic regions while retaining streamline counts reflective of the diffusion signal profile.

Along with the pitfalls of tractography, analysis of the tracts also pose a challenge. As previously mentioned, outlier streamlines pose a problem in both evaluating reliability of tracts and assessing quantitative metrics. A clear example was seen in the R OR, where outlier streamlines persist even after attempting to automatically identify and remove outlier streamlines. These streamlines, due to their similar trajectory with presumed true underlying anatomy, inflate the measures of Euclidean distance with the template. If such streamlines are removed, a clear reduction in the Euclidean distance is seen. Additionally, streamline counts are used to assess reliability of identified tracts. While it is more likely that different subjects may have differing tract densities, a single, healthy subject should exhibit minimal change across scanning sessions. From our U-shaped tract analysis, we see the lack of streamlines for some tracts across the different sessions of the intrasubject dataset. While we have identified these tracts as unreliable, further exploration is needed. Finally, the quantitative metrics analysed (eg. FA, MD) were derived from diffusion tensor measurements, which suffer from known problems in regions of streamline crossing and kissing. Our developed tool is not limited to diffusion tensor measurements and can include higher order or more advanced quantitative measures.

## Conclusions

In this work, we demonstrate the use of our tractography clustering tool and evaluate its ability to reliably identify and assess white matter pathways from whole-brain tractography using both a single-subject, multi-session dataset (MyConnectome) and a multi-subject, single-session dataset (HCP). Identified tracts within each of these datasets demonstrated good spatial overlap, with the exception of the R OR and R AF, which were noted to contain outlier streamlines and difficulty in tracking respectively. Quantitative, scalar metrics were shown to exhibit excellent reproducibility within each dataset. Additionally, 125 superficial white matter (U-shaped) tracts were determined to be reliable using the tractography generated across the two datasets.

In the future, we look to assess these identified U-shaped tracts to acquire a better understanding of their biophysical properties, their relation to cortical measurements and how these pathways are affected in patient populations.

## Acknowledgements

This research was enabled in part by the support provided by Compute Canada (www.computecanada.ca). Data was provided in part by the Human Connectome Project, WU-Minn Consortium (Principal Investigators: David Van Essen and Kamil Ugurbil; 1U54MH091657) funded by the 16 NIH Institutes and Centers that support the NIH Blueprint for Neuroscience Research; and by the McDonnell Center for Systems Neuroscience at Washington University.

